# Testosterone modulates status-specific patterns of cooperation in a social network

**DOI:** 10.1101/453548

**Authors:** T. Brandt Ryder, Roslyn Dakin, Ben J. Vernasco, Brian S. Evans, Brent M. Horton, Ignacio T. Moore

## Abstract

Stable cooperation requires plasticity whereby individuals are able to express competitive or cooperative behaviors depending on social context. To date, however, the physiological mechanisms that underlie behavioral variation in cooperative systems are poorly understood. We studied hormone-mediated behavior in the wire-tailed manakin (*Pipra filicauda*), a gregarious songbird whose cooperative partnerships are crucial for fitness. We used automated telemetry to monitor > 36,000 cooperative interactions among male manakins over three field seasons, and we examined how circulating testosterone affects cooperation using > 500 hormone samples. Observational data show that in non-territorial floater males, high testosterone is associated with increased cooperative behaviors and subsequent ascension to territorial status. In territory-holding males, however, both observational and experimental evidence demonstrate that high testosterone antagonizes cooperation. Moreover, circulating testosterone explains significant variation (2-8%) in social behavior within each status class. Collectively, our findings show that the hormonal control of cooperation depends on a male’s social status. We propose that the status-dependent reorganization of hormone-regulatory pathways can facilitate stable cooperative partnerships, and thus provide direct fitness benefits for males.

## Introduction

Behavioral plasticity is hypothesized to play a central role in the evolution of stable cooperation (Bergmuller et al. 2010; Díaz-Muñoz et al. 2014). For example, individuals must balance competition for mating opportunities with the benefits of engaging in cooperation (Clutton-Brock 2009; Cockburn 1998). These negotiations within cooperative social systems are mediated by physiological mechanisms that govern behavioral plasticity (Soares et al. 2010) over the course of different temporal scales. Over the long-term, the plasticity that drives among-individual differences in behavior and social status can facilitate stable cooperative partnerships (Bergmuller et al. 2010) as well as the competitive behaviors essential for establishing dominance and acquiring status within social groups (Ryder et al. 2008, 2009). Over shorter timeframes, within-individual plasticity is thought to facilitate the expression of cooperative or competitive behaviors that are appropriate for the current social context (Taborsky and Oliveira 2012). To date, however, the proximate underpinnings of the behaviors that promote stable cooperation are poorly understood (Soares et al. 2010). Thus, a key question is how physiological mechanisms modulate behavior in social systems where fitness requires a balance of cooperative and competitive behaviors.

Steroid hormones are well known for their pleiotropic and often antagonistic effects on numerous aspects of physiology and behavior (Ketterson et al. 1996; Wingfield et al. 2001; Adkins-Regan 2005; Hau 2007). Androgens, in particular, have been well studied in male vertebrates, where they can either facilitate or constrain behavioral plasticity in reproductive and social contexts (Adkins-Regan 2005; Hau 2007). Yet, the vast majority of work on androgen-mediated traits has focused on archetypal behaviors (e.g., aggression), with very few studies attempting to link hormones to individual variation in cooperation (Khan et al. 2001; Soares et al. 2010; Díaz-Muñoz et al. 2014). Although studies of cooperative social systems have documented status-specific differences in circulating testosterone levels (Wingfield et al. 1990; Schoech et al. 1991; Peters et al. 2001; Ryder et al. 2011*b*), including the suppression of androgens in subordinates (Brouwer et al. 2009), no study has examined how androgens mediate the dynamics of cooperation and status within a social network.

Circulating hormones have long been recognized as a key mechanism that may underlie individual differences in behavior. Mounting evidence suggests that hormonal phenotypes (i.e., the expression of consistently low or high hormone levels) are often repeatable and may thus act as a source of heritable variation driving individual differences in behavior (Williams 2008; Mills et al. 2009; While et al. 2010; van Oers et al. 2011; Iserbyt et al. 2015). However, empirical studies linking among-individual differences in circulating hormones to behavioral variation are challenging (Niemelä and Dingemanse 2018), in part, because hormone-mediated behaviors are regulated by complex signaling pathways (Adkins-Regan 2005; Ball and Balthazart 2008; Rosvall et al. 2012; Fuxjager et al. 2018). Indeed, the complexity of steroid-signaling pathways can produce nonlinear dose-dependent responses to circulating hormones, and these non-monotonic relationships may be key to understanding how hormones shape within and among-individual differences in behavior (Lema 2014).

In addition to the above challenges, both steroid hormones and behavior are intrinsically labile and responsive to the social environment. Steroids not only affect behavior, but they can also dynamically respond to it, and this social modulation of hormone-behavior relationships is thought to optimize behaviors to the current social landscape (Wingfield et al. 1990; Goymann 2009; Oliveira 2009; Goymann et al. 2019). Engaging in social interactions can rapidly alter circulating testosterone levels, androgen sensitivity in neural pathways, and the patterns of neural gene expression that influence motivation and shape status-specific behavioral phenotypes (Burmeister et al. 2007; Maruska and Fernald 2010; Fuxjager et al. 2010). Transitions in social status can likewise alter circulating hormones and their impact on behavioral neural circuits (Romeo et al. 2002; Maruska and Fernald 2010).

In an effort to expand our understanding of how hormones regulate social behavior, we asked how circulating testosterone modulates cooperative behavior in a lek-breeding bird, the wire-tailed manakin (*Pipra filicauda*; Fig. 1A). Wire-tailed manakins have a social system with two male status classes: territory-holders and floaters who do not yet hold a territory. Males from both classes form coalitions and perform a cooperative courtship display on territories at long-established lek sites (Heindl 2002). These coalitions are important for reproductive success, as territorial males with more display partners sire more offspring, and floaters with more partners have a higher probability of eventually inheriting a territory, which is a prerequisite for fitness (Ryder et al. 2008, 2009). Wire-tailed manakins do not exhibit overt aggression during display bouts, but instead competition maintains dominance hierarchies among coalition partners (Heindl 2002; Ryder et al. 2008). As such, cooperative coalitions have clear fitness benefits, but they also involve male-male competition and play a role in the ascension to territorial status. Many coalition partnerships are stable from year to year, but males also show substantial day-to-day variation in their social interactions, creating a dynamic social network (Ryder et al. 2011*a*; Dakin and Ryder 2018). Previous work on this system has established that the status classes differ in their circulating testosterone levels, with territory-holders having higher levels than floaters (Ryder et al. 2011*b*), and that cooperation is socially contagious (Dakin and Ryder 2018).

**Fig. 1.**
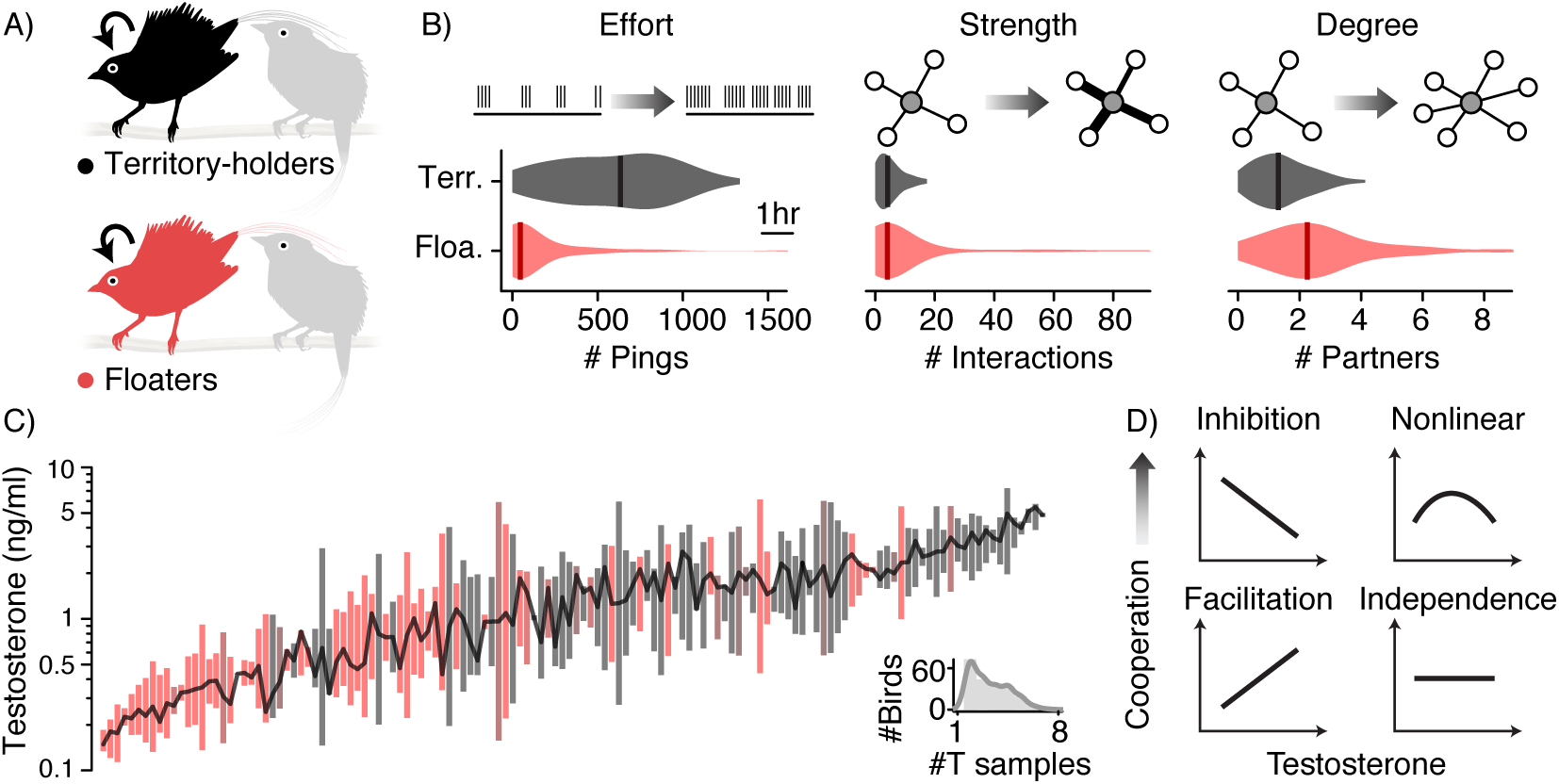
We studied how testosterone influences cooperative behavior in a social network. (A) Wire-tailed manakins have a social system in which males of two status classes, territory-holders (black) and floaters (red), form cooperative display coalitions. Once a male attains territorial status, he retains it for life. (B) We used an automated proximity system to quantify three social behaviors important for male fitness: effort is a measure of lek attendance; strength is the frequency of cooperative interactions; and degree is the number of unique cooperative partners; each was measured on a daily basis. The violin plots show the distributions of individual means (n = 180 individuals); the vertical lines are the medians. See Tables A1 and A2 in the Appendix for additional data. (C) We also collected repeated testosterone samples from 210 individuals (inset; n = 524 blood samples). Analysis of these repeated measures revealed that circulating testosterone (T) varies both among- and within-individuals, allowing us to partition these two sources of variation (Fig. A1 and Table A3 in the Appendix). The plot in (C) shows log-transformed testosterone for males with >1 hormone sample; each bar represents a single male, ordered along the x-axis by his average testosterone after correcting for capture conditions. Each individual’s bar spans the 25^th^ and 75^th^ quantiles and is colored according to his status; the dark line connects the medians. (D) We used these data to evaluates the predictions that testosterone may inhibit, facilitate, or have nonlinear (dose-dependent) effects on cooperation; the null hypothesis is that behavior is independent of circulating testosterone. We also considered that these hormone-behavior relationships may differ between the two status classes in (A).

Our goal in this study was to evaluate the role of a male’s circulating testosterone in shaping his behavioral phenotype (Fig. 1A). To quantify cooperative behaviors and identify coalition partners, we utilized an autonomous behavior-logging system to monitor the social activity and interactions of 180 males (Ryder et al. 2012; Dakin and Ryder 2018). We focused on three behavioral phenotypes measured for each male on a daily basis (Fig. 1B): “effort”, represents the duration of time a male spent attending the leks; “strength”, his frequency of cooperative interactions; and “degree”, his number of unique social partners. We consider effort to be a measure of status seeking and status maintenance for floater and territorial males, respectively, whereas strength and degree are two measures of a male’s cooperative tendencies. In addition to repeated measures of behavior, we also collected repeated hormone samples from 210 individuals in our study population to characterize circulating testosterone levels over time; each male was hormone sampled up to three times per field season (Fig. 1C). We used these hormone data to partition the standing variation in testosterone within and among-individuals, while controlling for capture conditions (Williams 2008; Vernasco et al. 2019), and examined the relationships between testosterone and behavior.

Our analysis evaluates three predictions for the hypothesis that testosterone shapes male behavioral phenotype (Fig. 1D). First, given that testosterone plays a well-known role promoting competitive ability and aggression in reproductive contexts (Goymann 2009), we consider that testosterone may be negatively related to cooperative behavior (i.e., “inhibition” in Fig. 1D). In contrast, evidence in humans suggests that testosterone may promote reciprocity and fair bargaining (Eisenegger et al. 2010; Boksem et al. 2013). As such, our second prediction was that testosterone may be positively related to cooperation (i.e., “facilitation” in Fig. 1D). Third, substantial evidence suggests that above some threshold, testosterone will no longer be correlated with behavior due to various mechanisms (Grunt and Young 1952; Kohn and Melnick 2002; Adkins-Regan 2005), or it may have detrimental effects on prosocial behaviors (e.g., parental care; Wingfield et al. 2001). As such, our third prediction was that testosterone may modulate cooperation in a nonlinear fashion (i.e., inverted U-function in Fig. 1D), whereby individuals with intermediate circulating levels are the most cooperative. Given that wire-tailed manakins have two status classes, and that social status is known to modulate hormone-behavior relationships in other species (e.g., Maruska and Fernald 2010), our analysis also tested for status-dependence in the relationship between testosterone and behavior. We also examined the null hypothesis that social behavior may be decoupled from testosterone (i.e., “independence” in Fig. 1D; Moore et al., 2004; Lynn et al., 2005; Hau, 2007). Finally, to establish a causal relationship between circulating testosterone and cooperation, we paired our observational analysis with a hormone manipulation experiment involving a subset of territorial males.

## Methods

### Study System

We studied a population of wire-tailed manakins (*Pipra filicauda*) that has been investigated and color-banded annually since 2002 at the Tiputini Biodiversity Station in Orellana Province, Ecuador (0° 38’ S, 76° 08’ W, approx. 200 m elevation). The breeding season lasts up to six months from November to April. Social status for each individual (i.e., territorial or floater) was determined each year by direct observation (Ryder et al. 2008). Briefly, if an individual was seen displaying on a territory repeatedly and consistently over several days or weeks, he was assigned as the territory owner. All blood samples and behavioral data were collected during peak breeding activity (December to March) over three field seasons (2015-16, 2016-17, and 2017-18). Sample sizes were not predetermined, but rather we tried to capture and sample all males at the studied leks each field season.

### Testosterone Sampling and Assay

Each male manakin was captured using mist-nets up to three times per field season. To maximize capture rates and minimize disturbance on the display territories (Vernasco et al. 2019), we deployed up to 16 mist-nets simultaneously on a given lek. Nets were checked every 30 minutes. A previous study demonstrated a subtle but significant effect net time (i.e., time from capture to blood sampling) on circulating testosterone in manakins (Vernasco et al. 2019). Therefore, we used video monitoring of the mist nets to account for net time in our subsequent statistical analyses (described below). The average span of time between repeated captures was 39 days (range 8-88 days). Blood samples (75-100uL) were collected from the brachial vein and stored on ice until return from the field when they were centrifuged for 5 min at 10,000 rpm. After centrifugation, plasma volume was measured to the nearest 0.25ul and stored in 0.75ml of 100% ethanol (Goymann et al. 2007); plasma volumes averaged 38.6uL (SD ± 10.3uL). In the lab, testosterone was double extracted from plasma with dichloromethane (Ryder et al. 2011*b*). We used direct radioimmunoassay to determine the total plasma testosterone concentration (ng/mL), adjusted by extraction efficiency and by the plasma volume of the sample (Eikenaar et al. 2011). Any sample that fell below the detection limit was assigned the assay-specific limit of detection as its testosterone concentration (0.12, 0.08, and 0.09 ng/mL for 2015-16, 2016-17 and 2017-18, respectively). Extraction efficiency ranged from 62-73% and the intra-assay coefficients of variation were 6.6%, 11.6% and 9.2% for 2015-16, 2016-17 and 2017-18, respectively. The inter-assay coefficient of variation was 19.5%.

### Partitioning Variation in Testosterone

A total of 524 testosterone samples were obtained from 210 individuals (1-8 samples per male; Fig. 1C). To evaluate whether testosterone differed among individuals (Williams 2008; Hau and Goymann 2015), we fit a mixed-effects model of log-transformed circulating testosterone in the lme4 package (Bates et al. 2018) in R (R Core Team 2018). The model included a bird’s ID as a random effect and controlled for capture conditions that can influence hormone levels by using fixed effects of field season (2015-16, 2016-17, or 2017-18), Julian date within the field season, time of day, and net time (i.e., see Vernasco et al., 2019). For 51 captures that were missing net time data, we assumed the seasonal average net time (16-19 min) to ensure that the missing values would not influence the analysis. We then computed the repeatability as the proportion of total variance in circulating testosterone explained by among-individual differences, after correcting for the fixed effects above. The 95% confidence intervals for repeatability were determined by parametric bootstrapping. To characterize the change in testosterone during social ascent, we added a fixed effect of four ascent categories to the same analysis (i.e., floaters who did not subsequently ascend, floaters who subsequently acquired a territory, new territory-holders, and established territory-holders). We then used post-hoc Tukey tests and the Benjamini-Hochberg procedure to compare status and ascent categories.

To examine hormone-behavior relationships, we used “corrected T”, which we define as the residuals from the linear model of log-transformed testosterone, after accounting for capture conditions (i.e., field season, Julian date, time of day, and net time). Next, we used within-subject centering to partition the variation in corrected T within- and among-individuals. The calculations followed the within-subject centering procedure in (van de Pol and Wright 2009); see Fig. A1 for a detailed illustration of this procedure. Briefly, we first computed each male’s “mean T” as the average of his corrected T values. Next, we computed “relative T” by subtracting each male’s mean T from his corrected T values. Hence, mean T is a variable that captures among-individual variation in testosterone (with one value of mean T per male). In contrast, relative T represents a measure of a male’s testosterone in a given blood sample, relative to his overall average (one value per blood sample).

### Social Behavior

Social behaviors (Fig. 1B) were quantified on the display territories using an automated proximity data-logging system (described in Ryder et al. 2012; Dakin and Ryder 2018). Males were outfitted with coded nano-tags (NTQB-2, Lotek Wireless; 0.35 g), each of which transmitted a unique VHF signal ping once per 20 s (range: 19.5-20.5 s). Approximately 10 (±SD 7) days after capture, a proximity data-logger (SRX-DL800, Lotek Wireless) was placed within each territory at that lek to record social behaviors within a detection radius of approximately 30 m (i.e., the average diameter of a territory; Ryder et al. 2012; Dakin and Ryder 2018). The data-loggers recorded all tag pings emitted by males within each territory from 0600 to 1600 local time for ∼6 consecutive days (SD ± 1 day). We attempted to recapture males at each lek for additional blood samples one to two more times per field season, after which social behaviors were again recorded using the procedure described above. Thus, behaviors were monitored after hormone sampling, and hence we examine the relationship between testosterone levels and subsequent behavior. We recorded a total of 29,760 hours of social activity (2015-16: 49 territories, mean = 16 recording days per territory; 2016-17: 54 territories, mean = 21 days per territory; 2017-18: 48 territories, mean = 22 days per territory, SD = ±4 days in all field seasons). The number of males tagged for behavioral monitoring was n_2015-16_ = 100, n_2016-17_ = 114, and n_2017-18_ = 82, for a total of n_tagged_ = 180 unique individuals. Note that the number tagged individuals is fewer than the number of blood-sampled males, because some males were captured at unmonitored leks and not tagged. These additional individuals were included in the hormone repeatability and variance partitioning analysis to obtain more accurate estimates of each effect, but they were not part of the behavioral study.

We defined three behavioral phenotypes. A male’s effort, or his rate of lek attendance, was quantified as his number of pings per day at monitored display territories. To define strength (frequency of cooperative interactions) and degree (number of unique social partners), the proximity data were first filtered to identify times when two males co-occurred on a display territory as an indication of unique cooperative interactions (Ryder et al. 2008, 2012; Dakin and Ryder 2018). A detailed description of the filtering algorithm is provided in (Dakin and Ryder 2018); a ground-truthing experiment in that study also confirmed that joint detections represent individuals that were on average < 5 m apart. This spatial range corresponds to the visual and acoustic contact required for a typical display interaction (Heindl 2002). Extensive behavioral observations of our study population have demonstrated that overt aggression (i.e., physical contact and chasing) is exceedingly rare on the display territories. As such, this automated approach provides a reliable measure of cooperative interactions and increases both the quantity and quality of social network data relative to other methods (Ryder et al. 2012). An additional validation study also confirmed that the social interactions defined by this automated system corresponded to cooperative display coalitions that were observed directly (Ryder et al. 2012). In total, we identified 36,885 unique social interactions over the present three-year study. Note that for floaters, our measures of effort, strength and degree included activity on any territory, whereas for territory-holders, the same measures were limited to a male’s activity on his own territory. Behavioral measures were transformed using log(x + 1) to meet the assumptions of subsequent statistical analyses, and all measures were significantly repeatable (Dakin and Ryder 2018).

### Hormone Manipulation Experiment

To test whether testosterone has direct effects on male behavior, we conducted a within-subjects hormone manipulation experiment on a subset of the territory-holding males. We chose the territorial status class for this experiment because their high site fidelity makes them readily available for post-manipulation behavioral monitoring. Implant surgeries were performed on 14 randomly-selected territorial males during the peak of the breeding activity in two field seasons, 2016-17 and 2017-18. Subjects received a subcutaneous 7mm length × 1.47mm internal diameter silastic implant sealed at both ends with ∼1mm of silicone. The implants were either filled with crystalline testosterone (T treatment, n = 9) or left empty (the control, n = 8). Three of the males who were manipulated in 2016-17 also received the opposite treatment in the subsequent field season (n = 17 implants in total). Implant surgeries were scheduled between data-logger recording sessions, such that each treatment male had a 6-day (SD ± 1 day) pre-implant behavioral monitoring period within four weeks prior to implantation, and another 6-day (SD ± 1 day) post-implant behavioral monitoring period within 10 days after implantation. We verified that the timing of pre- and post-implant behavioral monitoring did not differ between the two treatment groups (pre-monitoring, p = 0.98; post-monitoring, p = 0.60, Tukey’s post-hoc tests on linear mixed-effects models).

Blood samples were also acquired in the weeks after implantation (mean 13 days, IQR 9-15 days) for 17 implant surgeries to verify that the T treatment significantly increased circulating testosterone relative to pre-implant levels (p = 0.04; pre-implant least-squares mean from a mixed-effects model = 1.67 ng/mL [95% CI = 0.92, 3.02]; post-implant = 2.93 ng/mL [1.67, 5.15]). In contrast, the control implant did not significantly alter circulating testosterone (p = 0.15; pre-implant = 1.21 ng/mL [95% CI = 0.55, 2.66]; post-implant = 0.57 ng/mL [0.26, 1.25]). Note that although the least-squares mean was lower in the post-implant control group, it was still within the 95% CI of the pre-implant control group, indicating no significant change. We also verified that all post-manipulation measures were within the natural (i.e., biologically relevant) range (see Fig. 1C and (Ryder et al. 2011*b*)). The average post-implant testosterone level in the T treatment group corresponded to the 78^th^ percentile among unmanipulated territory-holding males.

### Statistical Analyses

Prior to analyzing hormone-behavior relationships in the observational data, we removed data from individuals with a single blood sample, because quantifying relative T requires repeated sampling (Appendix Fig. A1). The remaining measures of daily behavior were each assigned a relative T value using the bird’s most recent prior hormone sample within four weeks (median 12 days, IQR 7-18 days). This step filtered the data to 2,363 daily recordings of 125 unmanipulated males. Next, we averaged the daily behavioral measures assigned to each blood sample (i.e., one row per unique measure of relative T). The final sample size was 346 hormone-behavior measures from 125 individuals, including 57 floaters and 76 territory-holders. Note that 56 + 76 > 125, because some males were studied at both the floater and territory-holder stage.

To evaluate the hormone-behavior relationships in these data, we used a model selection procedure in the MuMIn package (Bartoń 2018) in R (R Core Team 2018). Specifically, we evaluated support for competing models representing the hypotheses shown in Fig. 1D and defined in Table A4. All candidate mixed-effects models included a male’s identity as a random effect, as well as fixed effects of field season, Julian date within the field season, temperature (daily averages acquired at the Tiputini Biodiversity Station), the number of days since capture, social status, and its interaction with testosterone (except in the null model, which did not include testosterone). After the model selection procedure, statistical significance was determined using the lmerTest package (Kuznetsova et al. 2018) in R. In each case, we also tested for status-dependence of the hormone-behavior relationships by determining whether the interaction between status and a given testosterone metric was statistically significant. This interaction tested whether the slope of the relationship differed between floater and territorial males. Given that we found significant status-dependence for all three behaviors, we next repeated the model selection procedure for each status class separately. We present p-values from the best-fit model for each status class in our main results (see Appendix for complete details on both sets of analyses). To determine the percent of behavioral variance explained by the testosterone metrics within each status class, we obtained R^2^ estimates using the r2glmm package (Jaeger 2017) in R (R Core Team 2018).

To analyze the effect of the hormone manipulation experiment on behavior, we fit mixed-effects models that accounted for a male’s identity as a random effect as well as fixed effects of field season, temperature, and implant (either pre-implant, control, or T treatment; n = 211 daily measures of 14 implanted individuals). We used post-hoc Tukey’s tests in the multcomp package (Hothorn et al. 2017) in R (R Core Team 2018) to compare the treatments, and corrected p-values using the step-up false discovery rate procedure in the multtest package (Pollard et al. 2018). We conducted a further analysis to determine if the effect of the T implant depended on a bird’s pre-implant hormone level. To do this, we compared the fit of models with, and without, the statistical interaction between a male’s corrected T and the effect of the implant. We used corrected T for this interaction because the small sample size in the experiment was not sufficient for variance-partitioning. The corrected T value came from the blood sample taken at the time of surgery.

## Results

Both the floaters and territory-holders expressed a broad range of testosterone levels (Fig. 2A, Table 1). The floaters who successfully acquired a territory had significantly higher levels of circulating testosterone, as compared to other floaters who did not ascend (p = 0.04, Fig. 2B). Importantly, testosterone levels were significantly repeatable in the population, with 19% of the standing variation in hormone levels attributed to differences among-individuals. Thus, repeatable differences in hormone levels have the potential to explain at least some of the among-individual differences in behavior.

**Table 1.** Descriptive statistics for metrics of circulating testosterone. Quartiles are provided because the raw testosterone data have a log-normal distribution1. The sample size is n = 524 blood samples from 210 males (247 floater samples, and 277 territory-holder samples).

| Testosterone metric | Status class | 25<sup>th</sup><br>percentile | 50<sup>th</sup><br>percentile | 75<sup>th</sup><br>percentile |
| --- | --- | --- | --- | --- |
| Testosterone (ng/ml) | Floater | 0.17 | 0.37 | 1.21 |
|  | Territory-holder | 0.63 | 1.62 | 2.56 |
| Corrected T (residual) | Floater | 0.23 | 0.48 | 1.61 |
|  | Territory-holder | 0.73 | 2.01 | 3.47 |
| Mean T | Floater | -1.34 | -0.65 | 0.17 |
|  | Territory-holder | 0.03 | 0.37 | 0.89 |
| Relative T | Floater | -0.59 | -0.03 | 0.54 |
|  | Territory-holder | -0.52 | 0.04 | 0.59 |

<sup>1</sup>See the figshare repository for all data values: https://figshare.com/s/d36c1a5c822e00abc581

**Fig. 2.**
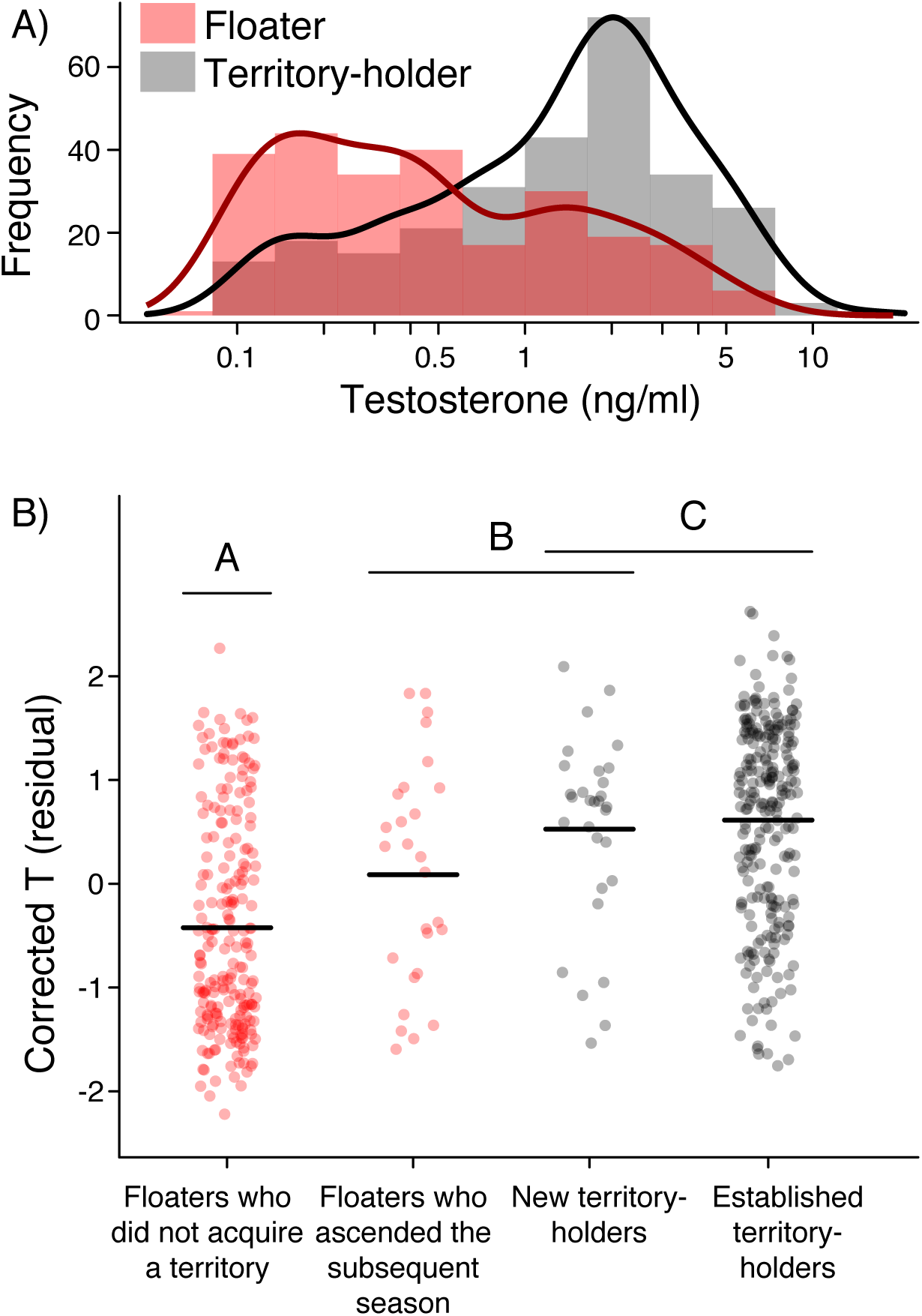
Elevated testosterone is associated with ascension to territorial status. (A) Histograms and probability density curves for log-transformed circulating testosterone in the two status classes (n = 524 samples). (B) Change in corrected T during ascension to territorial status. Floater males who acquired a territory the following field season had significantly higher testosterone than floaters who did not acquire a territory. The data points show partial residuals from an analysis that controls for field season and capture conditions (n = 524 measures from 210 individuals, including 15 who acquired territories during the study). Different letters indicate statistically significant Tukey contrasts, after correcting for multiple comparisons.

We found that testosterone could explain significant variation in all three behaviors, but these hormone-behavior relationships were highly status-specific, as evidenced by the significant slope differences between the two status classes (see Fig. 3). For floater males, the individuals with the highest levels of mean T attended the leks more often (effort) and engaged in cooperative interactions more frequently (strength; Fig. 3A-B). Floaters with higher mean T also tended to have more partners, although this relationship was not statistically significant (degree, p = 0.12; Fig. 3C). For territory-holding males, the males with higher mean T had significantly lower frequency of cooperative interactions (Fig. 3B), and tended to have fewer partners (p = 0.08; Fig. 3C). Lek attendance was highest in territory-holders with intermediate levels of mean T (Fig. 3A). In total, we found that the among-individual variation in testosterone (mean T) could explain between 2-8% of the variation (R^2^) in behavioral phenotypes within each status class (Fig. 3D). In contrast, the within-individual hormone variation (relative T) had a more limited role, and was primarily related to one behavior, lek attendance (effort), but only in territory-holders (Fig. A2 in the Appendix). The highest level of lek attendance was observed when a territory-holding male’s circulating testosterone was closest to his average. Together with the result in Fig. 3A, this indicates that territory-holders reduce their lek attendance above or below some threshold of mean T and relative T.

**Fig. 3.**
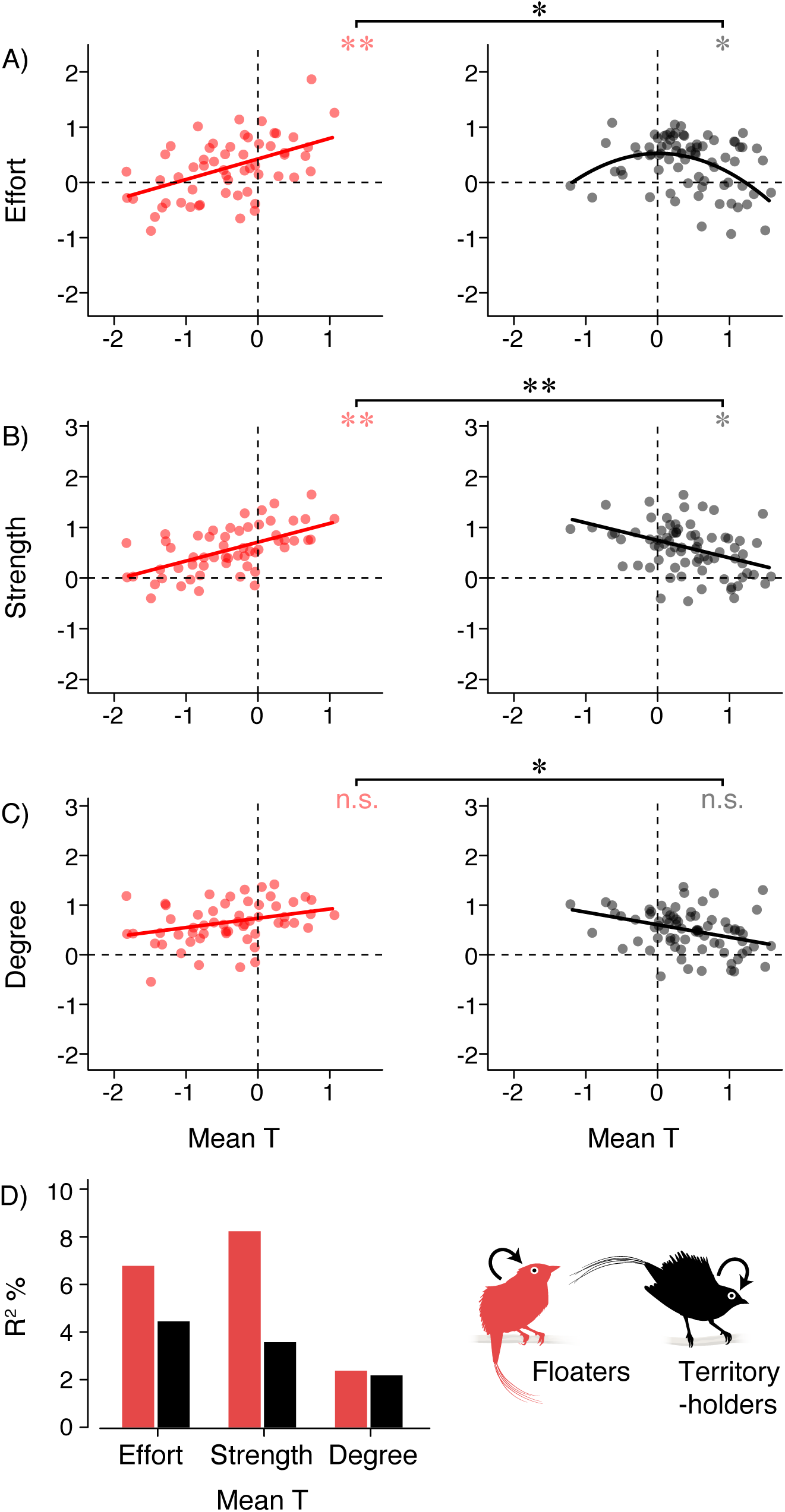
Social behavior is related to among-individual variation in circulating testosterone. (A-C) Mean T is a measure of among-individual variation in circulating testosterone. Floaters with the highest mean T had greater lek attendance (effort), cooperated with other males at higher frequencies (strength), and had more unique cooperative partners (degree), although the relationship with degree was not statistically significant (n = 57 floaters). In contrast, territory-holding males with the highest mean T had lower lek attendance, a lower frequency of cooperation, and fewer partners, although again the relationship with degree was not quite significant (n = 76 territory-holders). Each scatterplot shows the residuals (y-axis) from the best-fit model after accounting for other predictors; note that all phenotypes were analyzed on a log-scale. Red and grey asterisks indicate statistical significance within floaters and territory-holders, respectively; the black asterisks compare hormone-behavior relationships between the two status classes (n.s. non-significant; * p < 0.05; ** p < 0.01). (D) The percent of behavioral variance (R^2^) in each phenotype attributed to mean T. Further details are provided in Tables A4-A7 in the Appendix.

The territorial males with experimentally-elevated testosterone significantly decreased their cooperative behavior, both in terms of the number of display partners (degree), and the frequency of cooperative interactions (strength) they received, although the manipulation did not significantly affect their territory attendance (effort; Figure 4A-B). These experiment results are consistent with the correlative analysis (see Fig. 3), and confirm that testosterone levels above some threshold have a direct negative impact on a territorial male’s ability to maintain cooperative partnerships. Further analysis revealed that the effects of the T treatment also depended on a male’s pre-implant hormone level, as males with low testosterone prior to the implant lost more cooperative interactions and partners (strength and degree) than did males with initially high pre-implant testosterone (Fig. 4C).

**Fig. 4.**
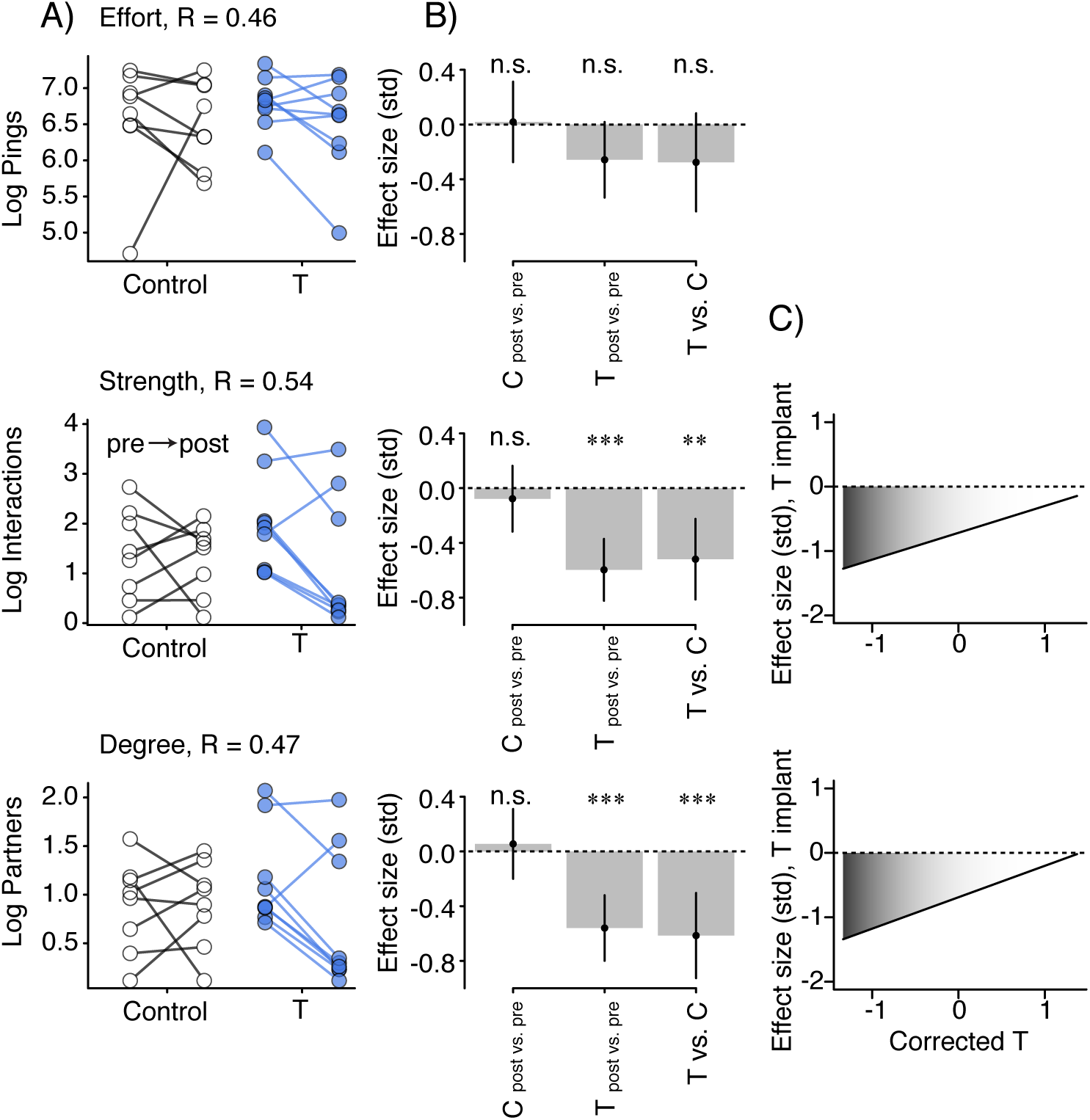
Experimental elevation of testosterone decreased cooperation in territory-holders. Territorial males were given control (n = 8) or testosterone (n = 9) implants. (A) Average pre- and post-manipulation measures of lek attendance (effort), the frequency of cooperative interactions (strength), and the number of partners (degree). Lines connect data from the same individual. The repeatability of each behavior, R, is given above each panel. (B) Standardized effect sizes from the statistical analysis (Control treatment, C post vs. pre; T treatment, T post vs. pre; and a comparison of the T treatment and the Control, T vs. C). Error bars show 95% confidence intervals. Experimentally-elevated testosterone decreased the frequency of cooperative interactions (strength) and the number of unique partners (degree), because T post vs. pre and T vs. C were both statistically significant after correcting for multiple comparisons (n.s. non-significant; * p < 0.05; ** p < 0.01; *** p < 0.001). The sample size for this analysis was n = 211 daily measures of 14 individuals. (C) Further analysis indicated that the effect of elevated testosterone on cooperative behavior (strength and degree) depended on the male’s pre-treatment corrected testosterone level. Specifically, stronger negative effects were observed for territory-holding males with lower pre-implant testosterone. See Tables A8-A9 in the Appendix for additional data. Standardized effects sizes are shown to be comparable across the figure.

## Discussion

The fitness of social animals is often contingent on the ability to express contextually relevant behaviors in response to a dynamic social environment. In the social system of wire-tailed manakins, floater males must cooperate with territory-holders and compete with other floaters for limited opportunities to acquire a territory (Ryder et al. 2008). However, once an individual ascends to territorial status, he must behaviorally shift his focus to attracting partners and building stable coalition partnerships on his own territory, because females prefer more cooperative individuals (Ryder et al. 2009). As such, selection is expected to favor endocrine mechanisms that promote behavioral plasticity to respond appropriately to conspecifics and ascension opportunities (Kelly and Vitousek 2017). Although we know that hormones can coordinate both within- and among-individual plasticity in behavior and social competence (Dufty et al. 2002; Hau and Goymann 2015), the vast majority work on androgens has focused on how they facilitate courtship, aggression, competitive abilities, and dominance (Ketterson et al. 1996; Wingfield et al. 2001; Fusani 2008).

Here, we show that testosterone, an androgen long-known for its effect on aggression, can also influence social behavior in a cooperative system. Higher average testosterone (mean T) was associated with an increased frequency of cooperation in floaters (Fig. 3). Therefore, it does not appear to antagonize cooperation, but instead may facilitate the social behaviors necessary for territory acquisition (Fig. 1D). Indeed, circulating testosterone was also predictive of which floaters ascended to territorial status, as the floaters that gained territories had higher testosterone than those who did not obtain territories, both pre- and post-ascension (Fig. 2). Given that territory ownership and coalition partnerships are both necessary for a male to sire offspring in this species (Ryder et al. 2008), these results suggest that high testosterone during the floater stage can yield later fitness benefits. Additional experiments are required to decouple the effects of age and testosterone on these correlative results, because the floaters that successfully ascend the age-graded status queue also tend to be older. Experiments that manipulate hormones or present social stimuli are also needed to determine whether higher testosterone causes status-seeking and cooperation in floaters, and/or whether testosterone rises in response to these behaviors.

Once a male acquires a territory, his fitness depends on both the number of cooperative partners and the stability of those partnerships (Ryder et al. 2009, 2011*a*). The results of our hormone manipulation experiment on territorial males confirmed that when testosterone in a territorial male is too high, it compromises his ability to attract and maintain stable display partnerships (Fig. 4). Hence, testosterone levels above some threshold at this stage could decrease reproductive fitness. Our correlative data also support this hypothesis, as territorial males with lower mean T exhibited the optimal combination of social behaviors (i.e., high effort, strength, and degree in Fig. 3). Therefore, testosterone levels appear to be ultimately constrained in territory-holders despite their potential benefits at the floater stage.

A key finding of our study is that the nature of hormone-behavior relationships depends on a male’s social status (Fig. 3), with all predictions in Fig. 1D represented, depending on the behavior in question and status of the individuals. This finding suggests that a reorganization of the hormone-regulatory pathways involving testosterone may be required to optimize status-specific social behaviors as males transition from floater to territorial. These changes may result from a number of physiological mechanisms that are not mutually exclusive. One possibility is that testosterone is only antagonistic to cooperative behavior above some concentration threshold that is not achieved by most floater males, and thus differential regulation of testosterone secretion according to social status is essential. The threshold at which testosterone inhibits cooperation may also depend, in part, on an individual’s hormonal phenotype or current endocrine status. This possibility is supported by the fact that the negative effect of T-implants on cooperative behavior was greater for territory-holders with lower pre-implant testosterone (Fig. 4C). Another possibility is that status-specific expression of steroidogenic enzymes and/or steroid receptors can cause among-individual and/or status-specific differences in the neural sensitivity to testosterone (Romeo et al. 2002; Burmeister et al. 2007; Rosvall et al. 2012; Horton et al. 2014). Finally, testosterone can also have downstream effects on the release or actions of other neurohormones known to influence social behavior, including steroid-sensitive neuropeptide systems (e.g., arginine vasotocin, and mesotocin; Kimura et al. 1999; De Vries and Panzica 2006; Kabelik et al. 2010), and these pathways may vary between status classes. Each of these hypotheses represents a promising direction for identifying how variation in the hormone signaling pathways facilitates cooperation in the manakin lineage. Future work examining status-specific neuroendocrine gene expression and its relationship with behavior in this species is a pivotal next step.

Our repeated hormone sampling efforts also revealed that individual males differ significantly in their circulating testosterone (corrected T; Fig. 1C). This finding of significant repeatability is consistent with growing evidence that hormonal phenotypes are often repeatable and heritable in other systems (Williams 2008; Mills et al. 2009; While et al. 2010; van Oers et al. 2011; Iserbyt et al. 2015), yet robust estimates from wild populations, like those presented here, are still exceedingly rare (Cox et al. 2016). Although the proportion of behavioral variation explained by among-individual variation in testosterone (mean T) was small at 2-8% (Fig. 3D), these R^2^ values are likely underestimates because of the measurement error associated with a limited number of repeat hormone samples (mean 2.5 blood samples per male, IQR 1-4; Fig. 1C). Moreover, a recent meta-analysis estimated that hormone levels generally account for only 2% of the variation in behavior, on average (Niemelä and Dingemanse 2018). Thus, the fact that among-individual differences in circulating hormones can account for as much as 8% of the variation in the behavior of a free-living animal is an important result (Ball and Balthazart, 2008).

The social transmission of behaviors is hypothesized to play an important role in the evolution of cooperation (Trivers 1971; Rankin and Taborsky 2009). Our previous work has established that manakin cooperative behaviors are indeed socially contagious, whereby the cooperative behaviors of one individual stimulate subsequent cooperation in social partners (Dakin and Ryder 2018). Here, we provide evidence that testosterone can influence the expression of socially-contagious behavior, in particular the frequency at which territory-holding males engage in cooperative interactions (Figs. 3-4). This raises the hypothesis for future study that variation in testosterone levels may influence the collective structure and assortment of the social network. Because most coalition partnerships involve a territory-holder and a floater (Ryder et al. 2011*a*), status-specific differences in how testosterone mediates behavior may promote a stable social network, by facilitating cooperation between males of different status classes.

A recently proposed framework suggests that aggression and sociality are inextricably linked and modulated by the same physiological and neural systems (Kelly and Vitousek 2017). In humans, the effects of testosterone depend on social status; testosterone can drive aggression when individuals are socially challenged, but drive prosocial behaviors in the absence of those challenges (Boksem et al. 2013). Moreover in other vertebrate taxa, changes in social status may often involve restructuring of the neural circuits and thus the way in which hormones mediate social behaviors (Romeo et al. 2002; Burmeister et al. 2007; Maruska and Fernald 2011; Maruska 2015; Fernald 2015). In parallel with these discoveries, we show here that both the optimal level of testosterone and the way it modulates behavior may depend on a male’s social status. Overall, our findings support the hypothesis that status-dependent effects of androgens on social behavior may be a general feature of cooperative social systems where fitness depends on behavioral plasticity.

## Acknowledgments

We thank Camilo Alfonso, David and Consuelo Romo, Kelly Swing, Diego Mosquera, and Gabriela Vinueza. This work was supported by the National Science Foundation (NSF) IOS 1353085, the Smithsonian Migratory Bird Center, the Global Change Center at Virginia Tech, Millersville University, and the Tiputini Biodiversity Station of the Universidad San Francisco de Quito. Matt Fuxjager provided helpful feedback on an earlier version of this manuscript.

## Author contributions

T.B.R., B.M.H., and I.T.M. designed the research

T.B.R., R.D., B.M.H., B.J.V., and I.T.M. performed the research

R.D., T.B.R., and B.S.E. analyzed the data

T.B.R. and R.D. wrote the manuscript

All authors edited the manuscript

## Competing interests

The authors declare no competing interests.

## Additional information

See the Appendix PDF for Figures A1-A2 and Tables A1-A9.

## Data Availability

All materials necessary to reproduce the results including data and R code are available for reviewers at the following link and will be made public upon acceptance: https://figshare.com/s/d36c1a5c822e00abc581

## Ethics

All methods were approved by the Smithsonian ACUC (protocols #12-23, 14-25, and 17-11) and the Ecuadorean Ministry of the Environment (MAE-DNB-CM-2015-0008).

## Notes

#### Summary of Updates

updated the analysis and edited the text according to input from peer reviewers

