## Appendix for "Testosterone modulates status-specific patterns of cooperation in a social network"

**Table A1. Behavioral phenotypes of male wire-tailed manakins.** Grand means for each status class were calculated from n = 4,669 daily measures of 180 control individuals. Behavioral phenotypes of territory-holding males were calculated using pings and interactions received on the male's own territory, whereas floater behaviors included any territory visited by the floater male. Note that for effort, one ping corresponds to approximately 0.33 minutes of lek attendance. Thus, the average effort of floater and territorial males is about 49 and 195 minutes per day, respectively.

| Behavioral phenotype | Floater mean (SE) | Territory-holder mean (SE) |
|---|---|---|
| Effort *(pings per day)* | 147 (24) | 586 (36) |
| Strength *(interactions per day)* | 10.1 (1.6) | 5.1 (0.5) |
| Degree *(partners per day)* | 2.6 (0.2) | 1.4 (0.1) |

**Table A2. Behavioral correlations among- and within-individuals.** Each cell gives the median posterior estimate of the correlation coefficient (ranging from –1 to 1), followed by the [*95% central range*], derived from a multivariate analysis of n = 4,669 daily measures of 180 control individuals. The analysis accounts for status, field season (fixed effects), and ID (random effect).

| | Effort | Strength | Degree |
|---|---|---|---|
| Effort | -- | 0.77 *[0.69, 0.84]* | 0.72 *[0.63, 0.80]* |
| Strength | | -- | 0.95 *[0.93, 0.97]* |
| Degree | | | -- |

| | Effort | Strength | Degree |
|---|---|---|---|
| Effort | -- | 0.66 *[0.64, 0.68]* | 0.63 *[0.61, 0.65]* |
| Strength | | -- | 0.90 *[0.89, 0.90]* |
| Degree | | | -- |

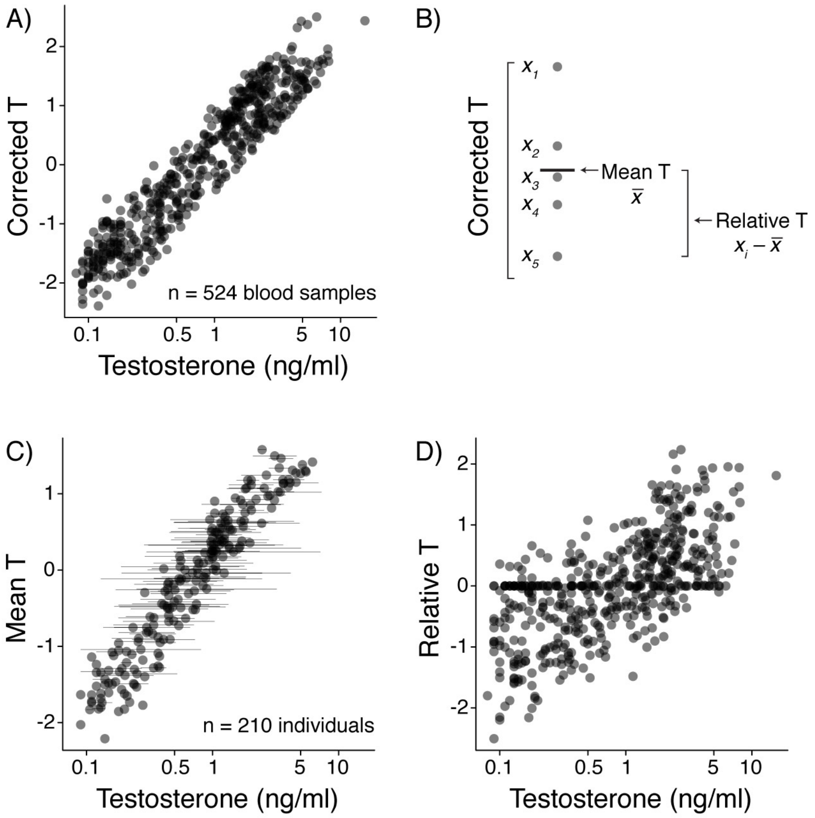


**Figure A1. The method of within-subject centering.** (A) Centering was performed on "corrected T", which is a residual value after statistically accounting for capture conditions. Here we show the correspondence between corrected T and the original testosterone concentrations. (B) Example of how centering was performed. The five data points represent five repeated measures of corrected T from the same individual, male *x*. "Mean T" for male *x* is the average of his five corrected T values. Each "relative T" value is calculated by subtracting mean T from one of the corrected T values. These two metrics, mean T and relative T, are orthogonal to each other, and provide a way to partition the among- and within-individual hormonal variation. (C-D) Correspondence between mean T, relative T, and the original testosterone concentrations. The scatterplot in (C) shows mean T in relation to a male's average uncorrected testosterone concentration (± SE; n = 210 values, one for each male). The scatterplot in (D) shows relative T in relation to uncorrected testosterone concentration (n = 524 values, one for each blood sample). Note that in (C), there are 76 data points where relative T = 0. These data points represent 76 individuals for whom only one blood sample was available. Although we have shown these 0 values here for illustration purposes, they were excluded from further hormone-behavior analyses because quantifying relative T requires >1 measure per individual.

**Table A3. Sources of variation in circulating testosterone**. The sample size is 524 blood samples from 210 males. Note that the response variable, circulating testosterone (ng/ml), was log transformed prior to analysis. Julian dates were assigned negative values in December counting backward from December 31=0; this created date variable that monotonically increased over each field season from December to March.

| Fixed effects | Estimate (SE) | t value | p-value |
|---|---|---|---|
| Field season | | | |
| 16-17 vs. 15-16 | –0.80 (0.12) | –6.88 | < 0.0001 |
| 17-18 vs. 15-16 | –0.41 (0.13) | –3.16 | 0.002 |
| Julian date | 0.001 (0.002) | 0.80 | 0.43 |
| Time of day (minutes) | –0.0006 (0.0004) | –1.36 | 0.17 |
| Duration in the net (minutes) | –0.018 (0.005) | –3.72 | 0.0002 |

| Repeatability analysis | Variance | Repeatability [95% CI] |
|---|---|---|
| Male ID | 0.27 | 0.19 [10, 30] |
| Residual | 1.11 | |

**Table A4. Candidate models of testosterone and behavior**. Seven candidate models are compared, with a bird's behavior as a function of his:

(1) mean T,
(2) relative T,
(3) mean T + relative T,
(4) mean T + a 2nd-degree polynomial function of relative T,
(5) a 2nd-degree polynomial of mean T + relative T,
(6) a 2nd-degree polynomial of both T variables, mean T and relative T, or
(7) (the null model), no testosterone variables.

All candidate models included the bird's identity as a random effect, as well as field season, Julian date, temperature, the number of days since capture, and status (territorial or floater) as fixed effects. Models (1)-(6) also included the interactions between the testosterone metrics and status; we tested this interaction to evaluate whether the hormone-behavior relationships differed significantly between the two status classes (i.e., status-dependence). Models are ranked by goodness-of-fit (AIC). The evidence ratios that compare the best-fit model to the null model for effort, strength, and degree are 10.0, 36.2, and 1.3, respectively.

| Behavioral phenotype | Model # | Candidate models | AIC | Delta | Akaike weight |
|---|---|---|---|---|---|
| Effort | 6 | poly(Mean T, 2)*Status + poly(Relative T, 2)*Status | 913.1 | 0 | 0.41 |
| | 4 | Mean T*Status + poly(Relative T, 2)*Status | 914.0 | 0.89 | 0.26 |
| | 1 | Mean T*Status | 916.2 | 3.05 | 0.09 |
| | 3 | Mean T*Status + Relative T*Status | 916.2 | 3.07 | 0.09 |
| | 5 | poly(Mean T, 2)*Status + Relative T*Status | 916.5 | 3.38 | 0.08 |
| | Null | No T variables | 917.7 | 4.59 | 0.04 |
| | 2 | Relative T*Status | 917.9 | 4.80 | 0.04 |
| Strength | 1 | Mean T*Status | 884.5 | 0 | 0.65 |
| | 4 | Mean T*Status + poly(Relative T, 2)*Status | 887.9 | 3.32 | 0.12 |
| | 3 | Mean T*Status + Relative T*Status | 888.4 | 3.87 | 0.09 |
| | 6 | poly(Mean T, 2)*Status + poly(Relative T, 2)*Status | 888.9 | 4.38 | 0.07 |
| | 5 | poly(Mean T, 2)*Status + Relative T*Status | 890.2 | 5.68 | 0.04 |
| | Null | No T variables | 891.7 | 7.21 | 0.02 |
| | 2 | Relative T*Status | 895.6 | 11.02 | < 0.01 |
| Degree | 1 | Mean T*Status | 847.7 | 0 | 0.46 |
| | Null | No T variables | 848.2 | 0.58 | 0.34 |
| | 3 | Mean T*Status + Relative T*Status | 851.6 | 3.98 | 0.06 |
| | 4 | Mean T*Status + poly(Relative T, 2)*Status | 852.1 | 4.47 | 0.05 |
| | 2 | Relative T*Status | 852.2 | 4.58 | 0.05 |
| | 5 | poly(Mean T, 2)*Status + Relative T*Status | 853.6 | 5.92 | 0.02 |
| | 6 | poly(Mean T, 2)*Status + poly(Relative T, 2)*Status | 853.5 | 5.95 | 0.02 |

**Table A5. Best-fit models for the associations between testosterone and behavior.** All phenotypic response variables, date variables, and daily temperatures were mean-centered and standardized (SD = 1). Each model also includes bird ID as a random effect. The sample size in each analysis is 346 measures of hormones and behavior from 125 control individuals.

| Behavioral phenotype | Best-fit model | | | |
|---|---|---|---|---|
| | Fixed effects | Estimate (SE) | t | p-value |
| Effort (log) | **Field season** | | | |
| | **16-17 vs. 15-16** | **0.74 (0.12)** | **5.99** | **< 0.0001** |
| | **17-18 vs. 15-16** | **0.68 (0.14)** | **4.93** | **< 0.0001** |
| | Julian date | –0.02 (0.05) | –0.42 | 0.68 |
| | **Mean temperature** | **0.11 (0.05)** | **2.05** | **0.04** |
| | Days since capture | 0.01 (0.05) | 0.23 | 0.82 |
| | **Status (Terr. vs. Floa.)** | **0.50 (0.16)** | **3.13** | **0.002** |
| | poly(Mean T, 1) * Status | | –0.78 | 0.43 |
| | poly(Mean T, 1), Floa. | 3.75 (2.48) | 1.51 | 0.13 |
| | poly(Mean T, 1), Terr. | 0.96 (2.75) | 0.35 | 0.73 |
| | poly(Mean T, 2) * Status | | –1.60 | 0.11 |
| | poly(Mean T, 2), Floa. | 0.05 (2.10) | 0.02 | 0.98 |
| | **poly(Mean T, 2), Terr.** | **–4.97 (2.30)** | **–2.16** | **0.04** |
| | poly(Relative T, 1) * Status | | –1.33 | 0.18 |
| | poly(Relative T, 1), Floa. | –0.08 (1.25) | –0.06 | 0.95 |
| | **poly(Relative T, 1), Terr.** | **–2.34 (1.12)** | **–2.09** | **0.04** |
| | **poly(Relative T, 2) * Status** | | **–2.07** | **0.04** |
| | poly(Relative T, 2), Floa. | 0.34 (1.30) | 0.27 | 0.79 |
| | **poly(Relative T, 2), Terr.** | **–3.45 (1.30)** | **–2.66** | **0.009** |
| Strength (log) | **Field season** | | | |
| | **16-17 vs. 15-16** | **1.04 (0.12)** | **8.73** | **< 0.0001** |
| | **17-18 vs. 15-16** | **0.38 (0.13)** | **2.86** | **0.005** |
| | Julian date | –0.08 (0.05) | –1.77 | 0.08 |
| | Mean temperature | 0.09 (0.05) | 1.81 | 0.07 |
| | Days since capture | 0.03 (0.05) | 0.63 | 0.53 |
| | **Status (Terr. vs. Floa.)** | **–0.46 (0.13)** | **–3.54** | **0.0005** |
| | **Mean T * Status** | | **–3.25** | **0.001** |
| | **Mean T, Floa.** | **0.34 (0.12)** | **2.79** | **0.006** |
| | Mean T, Terr. | –0.22 (0.12) | –1.73 | 0.09 |
| Degree (log) | **Field season** | | | |
| | **16-17 vs. 15-16** | **1.05 (0.11)** | **9.35** | **< 0.0001** |
| | **17-18 vs. 15-16** | **0.28 (0.13)** | **2.22** | **0.03** |
| | **Julian date** | **–0.11 (0.04)** | **–2.53** | **0.01** |
| | **Mean temperature** | **0.13 (0.05)** | **2.53** | **0.01** |
| | Days since capture | 0.01 (0.05) | 0.30 | 0.77 |
| | **Status (Terr. vs. Floa.)** | **–0.63 (0.13)** | **–4.99** | **< 0.0001** |
| | **Mean T * Status** | | **–2.11** | **0.04** |
| | Mean T, Floa. | –0.20 (0.12) | –1.61 | 0.11 |
| | Mean T, Terr. | 0.16 (0.12) | 1.31 | 0.19 |

**Table A6. Best-fit models when floaters and territory-holders were analyzed separately.** To verify that results were broadly consistent, we examined the candidate model set listed in Table A4 for each status class separately (142 hormone-behavior measures for the 57 floater males, and 204 measures for the 76 territory-holders, respectively). Note that 57 + 76 is greater than the total of 125 individuals from Table A5, because some males ascended in status during the study, and hence were present in both the floater and territory-holder subsets.

| Status class & behavioral | Best-fit model | | | |
|---|---|---|---|---|
| phenotype | Fixed effects | Estimate (SE) | t | p-value |
| FLOATER males | | | | |
| Effort (log) | **Field season** | | | |
| | **16-17 vs. 15-16** | **1.04 (0.19)** | **5.43** | **< 0.0001** |
| | **17-18 vs. 15-16** | **0.84 (0.23)** | **3.69** | **0.0004** |
| | Julian date | 0.03 (0.07) | 0.44 | 0.66 |
| | Mean temperature | 0.07 (0.08) | 0.86 | 0.39 |
| | Days since capture | 0.13 (0.08) | 1.59 | 0.12 |
| | **Mean T** | **0.37 (0.13)** | **2.78** | **0.007** |
| FLOATER males | | | | |
| Strength (log) | **Field season** | | | |
| | **16-17 vs. 15-16** | **1.36 (0.17)** | **8.02** | **< 0.0001** |
| | **17-18 vs. 15-16** | **0.65 (0.20)** | **3.17** | **0.002** |
| | Julian date | –0.07 (0.06) | –1.13 | 0.26 |
| | Mean temperature | 0.10 (0.07) | 1.43 | 0.16 |
| | Days since capture | 0.13 (0.07) | 1.83 | 0.07 |
| | **Mean T** | **0.37 (0.12)** | **3.00** | **0.004** |
| FLOATER males | | | | |
| Degree (log) | **Field season** | | | |
| | **16-17 vs. 15-16** | **1.46 (0.16)** | **8.83** | **< 0.0001** |
| | **17-18 vs. 15-16** | **0.47 (0.20)** | **2.40** | **0.02** |
| | Julian date | –0.09 (0.06) | –1.50 | 0.14 |
| | Mean temperature | 0.11 (0.07) | 1.64 | 0.10 |
| | Days since capture | 0.10 (0.07) | 1.53 | 0.13 |
| | Mean T | 0.19 (0.12) | 1.58 | 0.12 |
| TERRITORY-holder males | | | | |
| Effort (log) | **Field season** | | | |
| | **16-17 vs. 15-16** | **0.65 (0.17)** | **3.74** | **0.0003** |
| | **17-18 vs. 15-16** | **0.67 (0.19)** | **3.57** | **0.0005** |
| | Julian date | –0.05 (0.07) | –0.71 | 0.48 |
| | Mean temperature | 0.14 (0.08) | 1.74 | 0.09 |
| | Days since capture | –0.04 (0.07) | –0.55 | 0.58 |
| | poly(Mean T, 1) | –2.08 (1.15) | –1.81 | 0.07 |
| | **poly(Mean T, 2)** | **–2.09 (1.07)** | **–1.95** | **0.05** |
| | poly(Relative T, 1) | –1.57 (0.89) | –1.77 | 0.08 |
| | **poly(Relative T, 2)** | **–2.34 (0.98)** | **–2.38** | **0.02** |
| Strength (log) | **Field season** | | | |
| | **16-17 vs. 15-16** | **0.84 (0.17)** | **4.91** | **< 0.0001** |
| | 17-18 vs. 15-16 | 0.24 (0.19) | 1.27 | 0.21 |
| | Julian date | –0.11 (0.07) | –1.55 | 0.12 |
| | Mean temperature | 0.07 (0.08) | 0.89 | 0.38 |
| | Days since capture | –0.04 (0.07) | –0.57 | 0.57 |
| | **Mean T** | **–0.35 (0.14)** | **–2.47** | **0.02** |
| | poly(Relative T, 1) | –0.20 (0.89) | –0.22 | 0.82 |
| | **poly(Relative T, 2)** | **–1.93 (0.97)** | **–1.98** | **0.05** |
| Degree (log) | **Field season** | | | |
| | **16-17 vs. 15-16** | **0.84 (0.17)** | **4.90** | **< 0.0001** |
| | 17-18 vs. 15-16 | 0.21 (0.18) | 1.16 | 0.25 |
| | **Julian date** | **–0.13 (0.07)** | **–1.98** | **0.05** |

| | | | |
|---|---|---|---|
| Mean temperature | 0.10 (0.08) | 1.26 | 0.21 |
| Days since capture | –0.05 (0.07) | –0.81 | 0.42 |
| Mean T | –0.25 (0.14) | –1.79 | 0.08 |

**Table A7. Percent of the behavioral variance explained ($R^2$) by variation in circulating testosterone.** Results are computed separately for each status class using the results from Table A6 above.

| T metric | Behavioral phenotype | Status class | $R^2$ (percent variance explained) | [95% CI] |
|---|---|---|---|---|
| Mean T | Effort | Floater | 6.8 % | [1.1, 16.4] |
| | | Terr.-holder | 4.4 | [0.1, 15.7] |
| | Strength | Floater | 8.2 | [1.8, 18.3] |
| | | Terr.-holder | 3.6 | [0.3, 10.1] |
| | Degree | Floater | 2.4 | [0.0, 9.6] |
| | | Terr.-holder | 2.2 | [0.0, 7.8] |
| Relative T | Effort | Floater | -- | -- |
| | | Terr.-holder | 3.9 | [0.1, 14.7] |
| | Strength | Floater | -- | -- |
| | | Terr.-holder | 1.9 | [0.0, 9.7] |
| | Degree | Floater | -- | -- |
| | | Terr.-holder | -- | -- |

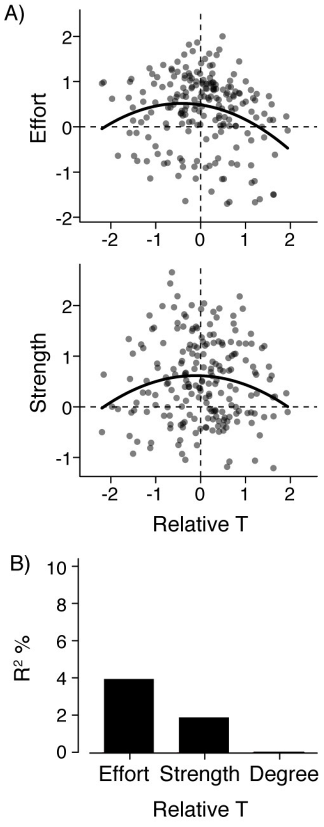


**Figure A2. Social behavior in relation to within-individual variation in circulating testosterone.** (A) Effort and strength were the only behavioral phenotypes significantly associated with relative T, and only within territory-holders (Table A6). Each scatterplot shows the residuals (y-axis) from the best-fit model after accounting for other predictors. The relationships between these two behavioral phenotypes and relative T are nonlinear. (B) The percent of behavioral variance ($R^2$) attributed to relative T in territory-holders.

**Table A8. Effects of experimentally-elevated testosterone on territorial males.** All phenotypic response variables and continuous predictors were mean-centered and standardized. Each model also includes bird ID as a random effect. We consider the effect of experimentally-elevated testosterone to be statistically significant if two criteria are met: (i) behavior changed significantly after the testosterone implant ("T implant post vs. pre"), and (ii) this change was significantly greater than that observed in the Control group ("T implant vs. Control"). The latter provides a control for the effects of capture and surgery. P-values from post-hoc Tukey's tests are reported after making a global correction for multiple comparisons. The sample size is 211 measures of 14 implanted individuals (105 pre- and 106 post-implant behavior measures).

| Behavioral phenotype | Fixed effects | Estimate (SE) | Test statistic | p |
|---|---|---|---|---|
| Effort (log) | Field season 17-18 vs. 16-17 | 0.53 (0.17) | 3.11 | |
| | Mean temperature | 0.16 (0.06) | 2.81 | |
| | Treatment effects | | | |
| | Control post vs. pre | 0.02 (0.15) | 0.13 | 0.99 |
| | T implant post vs. pre | –0.26 (0.14) | –1.83 | 0.16 |
| | T implant vs. Control | –0.28 (0.18) | –1.51 | 0.28 |
| Strength (log) | Field season 17-18 vs. 16-17 | –0.80 (0.14) | –5.67 | |
| | Mean temperature | 0.17 (0.05) | 3.72 | |
| | Treatment effects | | | |
| | Control post vs. pre | –0.08 (0.12) | –0.63 | 0.80 |
| | **T implant post vs. pre** | **–0.60 (0.12)** | **–5.18** | **< 0.0001** |
| | **T implant vs. Control** | **–0.52 (0.15)** | **–3.46** | **0.001** |
| Degree (log) | Field season 17-18 vs. 16-17 | –0.66 (0.15) | –4.46 | |
| | Mean temperature | 0.16 (0.05) | 3.18 | |
| | Treatment effects | | | |
| | Control post vs. pre | 0.05 (0.13) | 0.42 | 0.90 |
| | **T implant post vs. pre** | **–0.56 (0.12)** | **–4.58** | **< 0.0001** |
| | **T implant vs. Control** | **–0.61 (0.16)** | **–3.87** | **0.0003** |

**Table A9. Pre-implant testosterone level (corrected T) modulates the effect of the testosterone implants.** This analysis evaluates the potential interaction between a male's initial hormone level (corrected T) and the effect of experimentally-elevated testosterone on his behavior. Corrected T was measured from the blood sample taken immediately before the implant surgery. For each behavioral phenotype, goodness-of-fit was compared for the model with, and without, the interaction. ΔAIC is the change in Akaike's information criterion after dropping the interaction term; positive values indicate a better model fit when the interaction was included. A significant p-value indicates that the effect of the implant depended on a male's initial hormone level. The sample size is 113 measures of 9 T-implanted individuals (55 pre-implant and 58 post-implant behavioral measures).

| Behavioral phenotype | Corrected T * Implant interaction | | |
|---|---|---|---|
| | t | ΔAIC | p |
| Effort (log) | 0.83 | –2.27 | 0.94 |
| Strength (log) | **4.10** | **13.36** | **0.002** |
| Degree (log) | **4.34** | **14.61** | **0.001** |
